# Characterisation of *Plasmodium vivax* lactate dehydrogenase dynamics in *P. vivax* infections

**DOI:** 10.1101/2023.06.12.544683

**Authors:** Pengxing Cao, Steven Kho, Matthew J. Grigg, Bridget E. Barber, Kim A. Piera, Timothy William, Jeanne R. Poespoprodjo, Ihn Kyung Jang, Julie A. Simpson, James M. McCaw, Nicholas M. Anstey, James S. McCarthy, Sumudu Britton

## Abstract

*Plasmodium vivax* lactate dehydrogenase (PvLDH) is an essential enzyme in the glycolytic pathway of *Plasmodium vivax*. It can also be used as a diagnostic biomarker. Quantitation of plasma PvLDH has been used as a measure of *P. vivax* biomass in clinical studies of uncomplicated and severe vivax malaria. With the increasing importance of PvLDH in studying *P. vivax* diagnosis and infection, improved characterisation of the dynamics of this biomarker is important. In this study, we developed mathematical models that capture parasite and matrix PvLDH dynamics in *ex vivo* culture and the human host. We estimated the biological parameters using *ex vivo* and *in vivo* longitudinal data of parasitemia and PvLDH concentration collected from *P. vivax*-infected humans using Bayesian hierarchical inference. We found that the *ex vivo* and *in vivo* estimates of PvLDH in a parasitized red blood cell differed significantly across the asexual life cycle, with *in vivo* estimates at least ten-fold higher than *ex vivo* estimates (for example, the median estimate of intraerythrocytic PvLDH mass at the end of the life cycle was 9.4×10^−3^ ng *in vivo* vs. 5.1×10^−4^ ng *ex vivo*). We also estimated the *ex vivo* PvLDH half-life to be 65.3 h (95% credible interval: 60.8—70.7 h), which is approximately three times longer than the median estimate of the *in vivo* PvLDH half-life, 21.9 h (16.7—29.9 h). Our findings provide an important foundation to further improve quantitative understanding of *P. vivax* biology and facilitate the development of PvLDH-based diagnostic tools.

## Introduction

Malaria is a vector-borne disease, with 247 million malaria cases and 619,000 deaths estimated worldwide in 2021 ^1^. *Plasmodium vivax* is the most geographically widespread species causing malaria, putting billions of people at risk of infection ^2,3^. Although diagnosis of malaria by light microscopy remains the primary method, rapid diagnostic tests (RDTs) that detect *Plasmodium*-specific biomarkers present in the peripheral blood are widely used. *Plasmodium* lactate dehydrogenase (pLDH) is an essential enzyme in the glycolytic pathway of malaria parasites, and was one of the first parasite biomarkers identified in the blood of *Plasmodium-*infected patients ^4,5^. The development of RDTs that use *P. vivax*-specific monoclonal antibodies to target pLDH (PvLDH) has enabled species-specific detection of vivax infection without microscopy in areas co-endemic for *P. falciparum* and *P. vivax* ^6^. Quantification of plasma PvLDH has also been used as a measure of *P. vivax* biomass in clinical studies of uncomplicated and severe vivax malaria ^7,8^.

An understanding of the quantitative dynamics of a given biomarker facilitates its application as a diagnostic tool ^9^ and may reveal novel applications. Compared to the *P. falciparum*-specific biomarker, histidine-rich protein-2 (HRP2), the dynamics of which have been extensively studied and modelled ^9–12^, the dynamics of PvLDH remains poorly characterised. No mathematical methods for quantifying the biological parameters governing PvLDH dynamics based on *in vivo* data are available. There is an increasing amount of PvLDH data collected from vivax malaria patients, and among participants in human challenge studies where parasite kinetics are measured in a controlled setting. These data enable the characterisation of both *ex vivo* and *in vivo* PvLDH dynamics using mathematical modelling, and allow us to infer estimates for biological parameters that are difficult to measure by direct sampling. In particular, recent human studies have confirmed the accumulation of *P. vivax* parasites in reticulocyte-rich tissues outside the circulation, such as the spleen ^13,14^, and to a lesser extent the bone marrow ^15,16^, with the splenic reservoir accounting for more than 98% of total-body *P. vivax* biomass in chronic infections ^13^. In the absence of invasive tissue sampling, quantification of any parasite biomass that is outside peripheral circulation remains an ongoing challenge, and therefore, the biomarker PvLDH could provide new ways to quantify non-circulating parasite populations.

In this study, we quantified the *ex vivo* and *in vivo* dynamics of PvLDH by fitting two mathematical models to experimental data. One model that captured the *ex vivo* dynamics of both parasites and PvLDH (referred to as the *ex vivo* model) was fitted to a set of longitudinal measurements of parasite count and PvLDH concentration in two short-term *ex vivo* parasite cultures conducted using *P. vivax* isolates from two Indonesian patients with *P. vivax* monoinfection. The second model capturing *in vivo* parasite levels and PvLDH dynamics (referred to as the within-host model) was fitted to longitudinal data of parasitemia and whole blood PvLDH concentration from eight adults experimentally-infected with *P. vivax* in a volunteer infection study (VIS). We report the *ex vivo* and *in vivo* estimates of two fundamental parameters that characterise PvLDH dynamics. Firstly, intraerythrocytic PvLDH, which describes the average net accumulation of PvLDH mass inside a parasitized red blood cell (pRBC) over an asexual life cycle and is a function of parasite age. Secondly, the PvLDH decay rate or half-life, which quantifies how quickly PvLDH decays after release from ruptured pRBCs.

## Results

### Quantification of *ex vivo* PvLDH dynamics

The *ex vivo* model was fitted to both parasite count data derived by microscopy and ELISA-based PvLDH measurements collected in two 54-hour *ex vivo* cultures of *P. vivax* from two Indonesian patients using Bayesian hierarchical inference (see Materials and Methods for further details about the model and the *ex vivo* experiments). PvLDH levels in culture media measured at six-hour intervals indicate the amount of PvLDH released by ruptured pRBCs, while PvLDH levels in red blood cell (RBC) pellets indicate the intracellular PvLDH accumulated in the pRBCs. Figs. 1A-1D present the results of the model fitting, and show that the *ex vivo* model captures the trends of the measured data, as visualised by the median and 95% prediction interval (PI) of parasite count and PvLDH levels over time. We observed a rapid increase in PvLDH in culture media immediately after 0h in patient 2 (as opposed to the gradual increase in media PvLDH for Patient 1); this could be explained by the presence and rupturing of a higher fraction of schizonts at the start of the culture in patient 2 compared to patient 1 (2% vs. 0% by microscopy; 5.5% vs. 0.1% by the median model prediction). Data points in Figs. 1E and 1F present the proportion of parasites at each stage of the asexual cycle in the culture as determined by microscopy. At the start of culture, microscopy indicated that parasites were 91% and 93% ring stages (in patient 1 and 2, respectively); these matured into >90% trophozoites by 14—17 hours, and finally into schizonts (41% and 30% of parasites in patient 1 and 2, respectively) by the end of the culture. The fitted model was able to predict the fraction of parasites at various asexual developmental stages, and the predicted time series (visualised by the 95% PIs with median prediction curves shown in Figs. 1E and 1F) were largely consistent with experimental measurements, further supporting the ability of the *ex vivo* model to quantitatively capture the dynamics of the *ex vivo* system.

**Figure 1:**
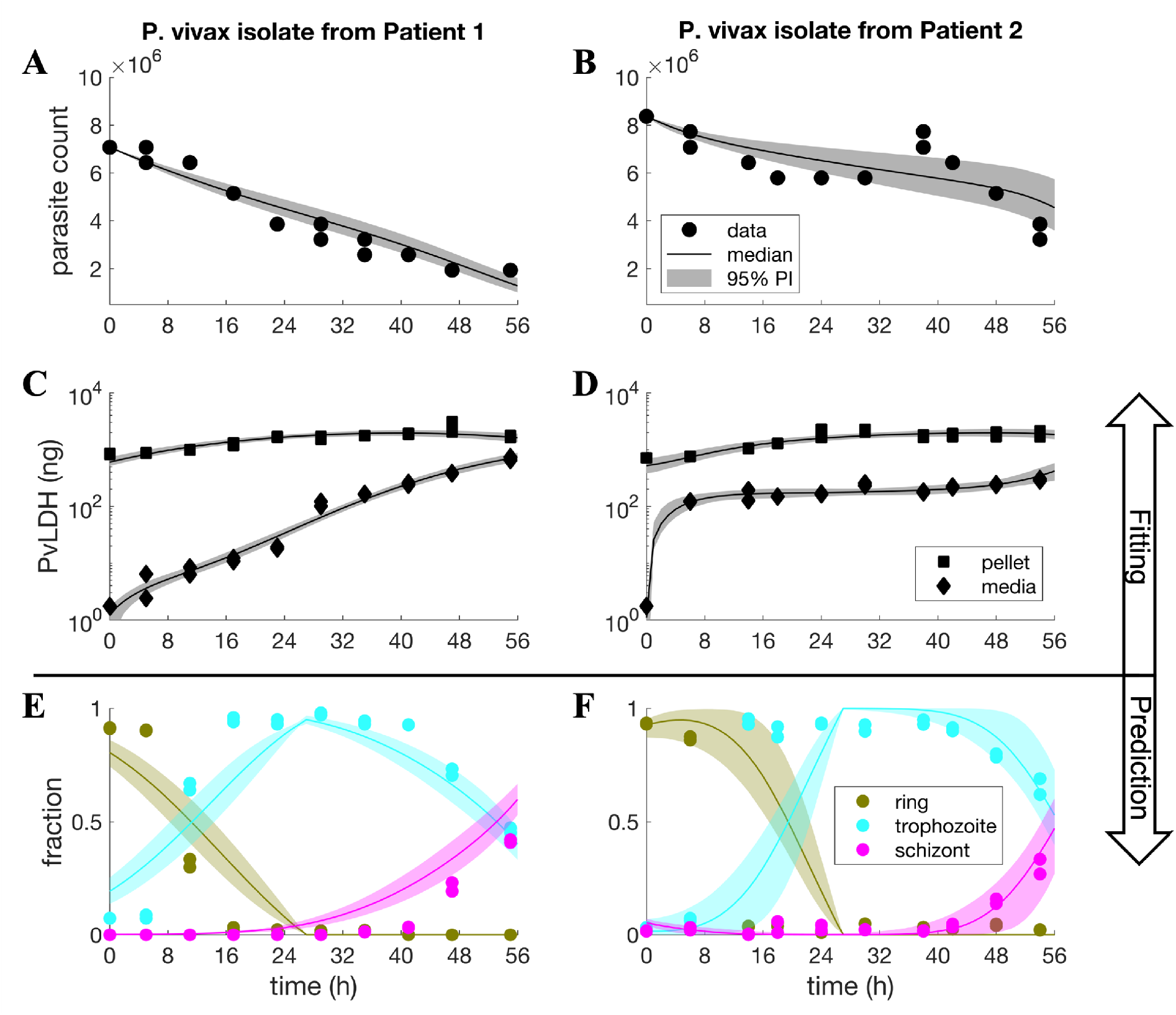
Results of *ex vivo* model fitting and model predictions. Panel A—D show the results of fitting the *ex vivo* model to the culture data of *P. vivax* isolates from two malaria patients in Timika, Indonesia. The solid dots are experimental measurements of parasite counts in the cultures and the solid squares and diamonds are experimental measurements of PvLDH in the culture pellet and media, respectively (note that PvLDH concentrations were measured and then converted to mass based on the volume of the culture; see Materials and Methods for details). Model fits are shown by the median and 95% prediction interval (PI) of the model-predicted distributions of parasite count and PvLDH levels (in nanogram) at different time points (in hours). The calculation of the median and the 95% PI is described in the Materials and Methods. Panel E and F show the model-predicted fractions of parasites at different stages of the lifecycle, which are shown by the median predictions (colour curves) and associated 95% PIs. Experimental measurements of the fractions of rings, trophozoites and schizonts are also shown in the figures for comparison (solid-coloured dots). Experimental measurements were conducted in duplicate at each timepoint.

Using the posterior samples of the population mean parameters obtained from the model fitting (marginal distributions of the samples are shown in Fig. S1 in the Supplementary Information), we estimated the *ex vivo* age-dependent accumulation of intraerythrocytic PvLDH and *ex vivo* PvLDH decay rate. Data modelling using a Hill function (Eq. 6 in the Materials and Methods) indicated that the median level of intraerythrocytic PvLDH was 2 to 3 orders of magnitude higher in mature stages (late trophozoite and schizont) compared to early stages (early and intermediate rings) (Fig. 2A). This is consistent with earlier findings that LDH levels are higher in blood samples where mature parasites are present ^4^. The maximum PvLDH per parasite at the end of the asexual life cycle was 5.1 ×10^−4^ ng (95% PI: 1.8 ×10^−4^—2.7 ×10^−3^) (Fig. 2A). The observed increase in PvLDH level as parasites age, suggests an accumulation of PvLDH produced by the parasite within the infected erythrocyte, and is in keeping with the slow increase in pellet PvLDH observed in the first 20—30 hours of the cultures (Figs. 1C and D).

**Figure 2:**
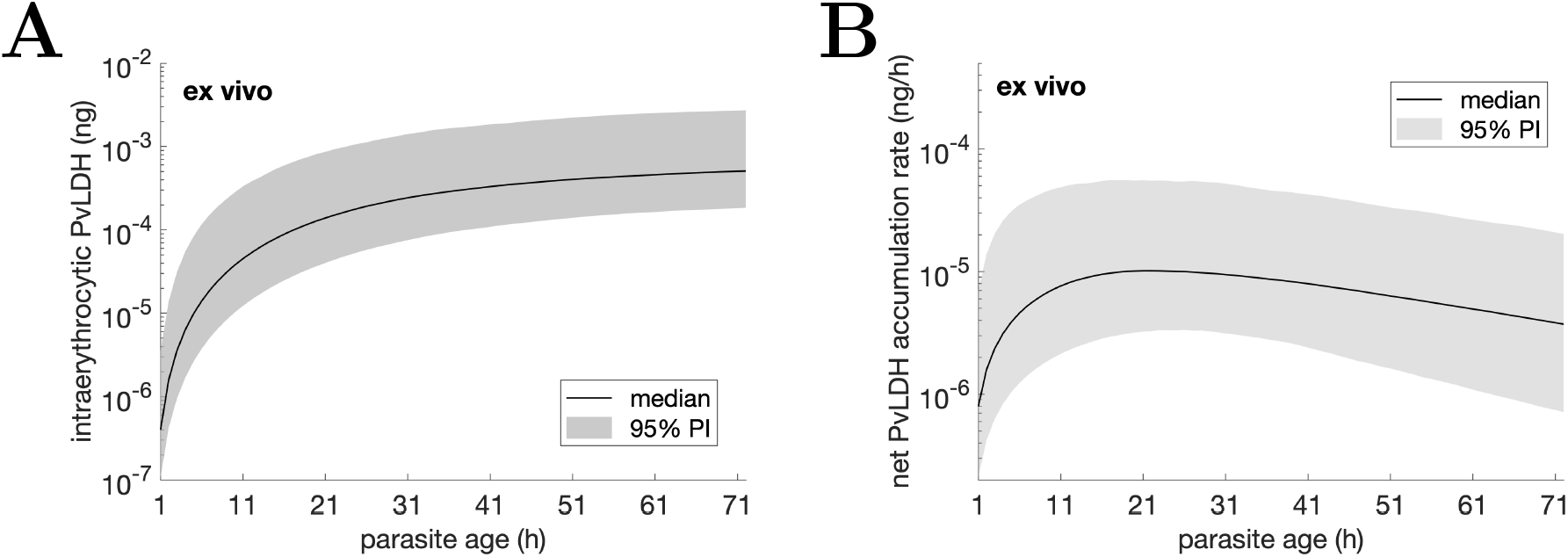
Estimated level of *ex vivo* intraerythrocytic PvLDH (ng) and net intraerythrocytic PvLDH accumulation rate (ng/h) over an *ex vivo* asexual life cycle. Note that the length of the *ex vivo* life cycle was estimated to be 72h (see Materials and Methods for the estimation of the *ex vivo* life cycle). The net intraerythrocytic PvLDH accumulation rate shown in B is the rate-of-change (the derivative) of the *ex vivo* intraerythrocytic PvLDH shown in A. Details of the calculation of the two quantities and their median and 95% prediction interval (PI) based on the posterior samples of population mean parameters are provided in the Materials and Methods.

The *ex vivo* PvLDH decay rate indicates the speed of decay of PvLDH in the culture media. Based on the marginal posterior distribution of the population mean of the *ex vivo* PvLDH decay rate (shown in Fig. S1 in the Supplementary Information), the *ex vivo* PvLDH decay rate was estimated to be 0.0106 h^-1^ (equivalent to a half-life of 65.3 h) and a 95% credible interval (CrI) of 0.0098—0.0114 h^-1^ (60.8—70.7 h). We also present the net PvLDH accumulation rate, calculated from the derivative of the intraerythrocytic PvLDH concentration, an indicator of the rate of accumulation of PvLDH inside the infected red cell (as a net result of production partially offset by degradation) (Fig. 2B). The *ex vivo* model predicts that the net rate of intraerythrocytic PvLDH accumulation peaks at approximately 20h from the start of an asexual life cycle during which parasites are at the ring stage (Fig. 2B).

### Quantification of *in vivo* PvLDH dynamics

Based on a similar framework used to construct the *ex vivo* model, we developed a within-host model to capture the *in vivo* dynamics of parasite growth and PvLDH turnover, and fitted the model to the parasitemia and whole blood PvLDH concentration data from eight human volunteers experimentally-infected with *P. vivax* (the isolate was originally from a patient with *P. vivax* malaria acquired in India) in a VIS (details about the VIS, the model and fitting method are provided in Materials and Methods). Figs. 3A—3D show the model fit and experimental data for two representative volunteers (R009 and R010; results for all eight volunteers are shown in Fig. S2 in the Supplementary Information). The model reproduced the synchronous stepwise growth pattern of *P. vivax* parasitemia and PvLDH production during the pre-treatment phase, as well as the decline in parasitemia and PvLDH levels after chloroquine treatment on day 10 (Figs. 3A—3D). We also used the model to predict the fraction of rings, trophozoites and schizonts over the period of observation, similar to the *ex vivo* study (albeit with no data for validation) (Figs. 3E and 3F). Consistent with the well-established life-cycle dynamics of *Plasmodium* infection, we found that the modelled distribution of parasite numbers across the life-cycle stages oscillates, and that the fraction of rings rapidly increases at the same time as a stepwise increase in parasitemia during the pre-treatment phase, timing that is consistent with schizont rupture.

**Figure 3:**
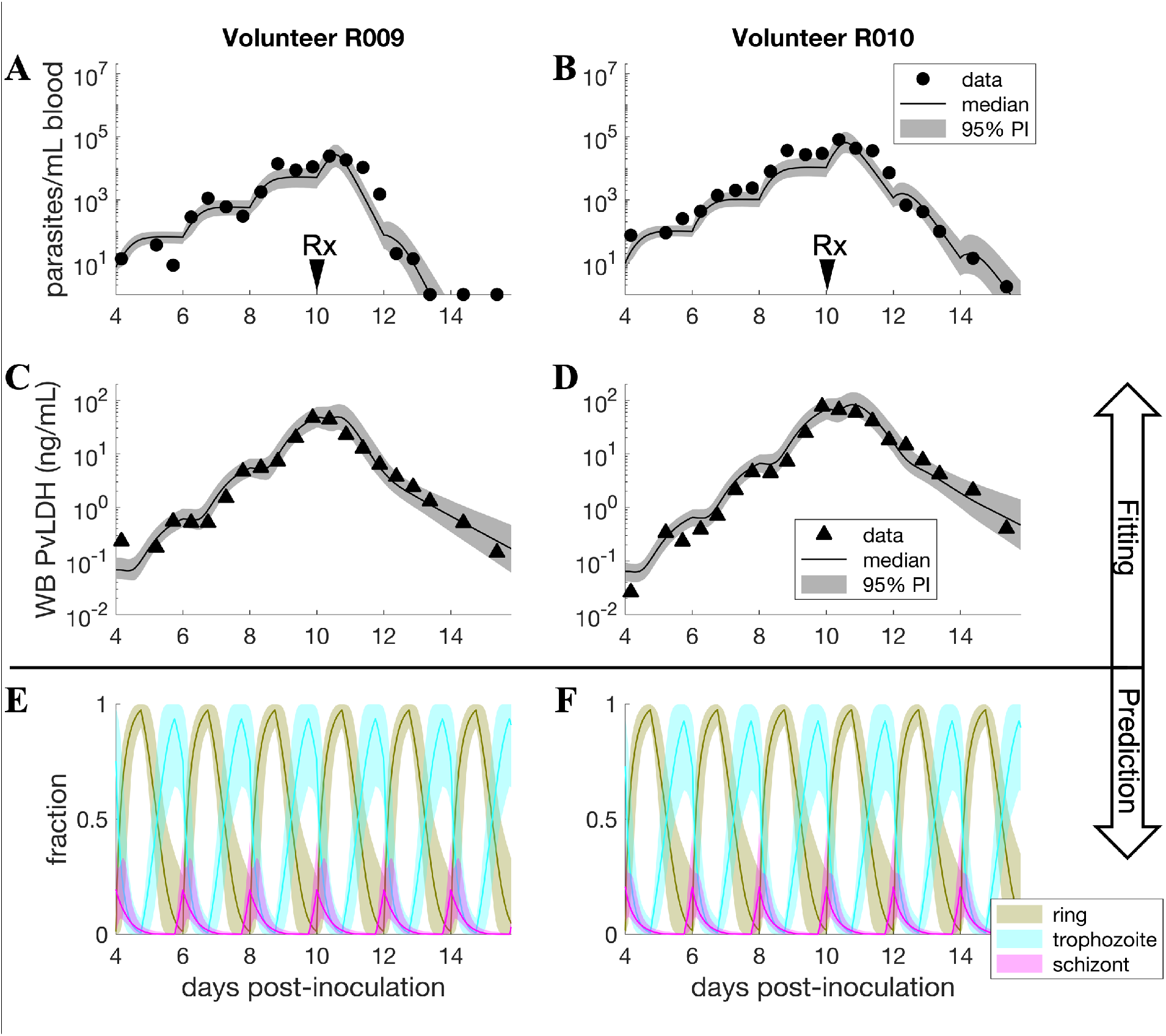
Results of within-host model fitting and predictions in VIS. Panel A—D show representative plots from two individuals after fitting the within-host model to parasitemia and whole blood (WB) PvLDH experimental data obtained from eight human volunteers experimentally-infected with *P. vivax* in a VIS. Model fits are shown by the median and 95% prediction interval (PI) of model-predicted distributions of parasitemia and WB PvLDH concentration at different time points. The calculation of the median and 95% PI is described in the Materials and Methods. The two volunteers were treated with a standard course of oral chloroquine on day 10 post-inoculation (indicated by the arrows labelled with Rx). Panel E and F show the model-predicted fractions of asexual developmental stages, illustrated by the median predictions (colour curves) and associated 95% PIs. For volunteer R009, some post-treatment parasitemia data are truncated at the lower detection limit of 1 parasite/mL.

We next calculated the *in vivo* intraerythrocytic PvLDH level using data from posterior distributions (for further details see Eq. 18 in the Materials and Methods and Fig. S3 in the Supplementary Information for the marginal posterior distributions of population mean parameters). The estimated *in vivo* intraerythrocytic PvLDH level reached a median value of 9.2 ×10^−3^ (95% PI: 2.1 ×10^−3^ —1.4 ×10^−2^) ng PvLDH per parasite at the end of the lifecycle (Fig. 4A), a figure that was approximately 18 times higher than the median of 5.1 ×10^−4^ ng per parasite estimated by the *ex vivo* model (Fig. 2A). Furthermore, we estimated the *ex vivo* and *in vivo* model-predicted amounts of intraerythrocytic PvLDH for each asexual developmental stage (Fig. 5). We found that the estimates of the intraerythrocytic PvLDH level in the *P. vivax* parasites across all asexual developmental stages in the VIS were at least ten-fold higher than the estimates of the intraerythrocytic PvLDH levels in the *ex vivo* cultures. In addition, the *in vivo* net intraerythrocytic PvLDH accumulation rate (Fig. 4B) was approximately one order of magnitude higher than the accumulation rate in *ex vivo* cultures (Fig. 2B).

**Figure 4:**
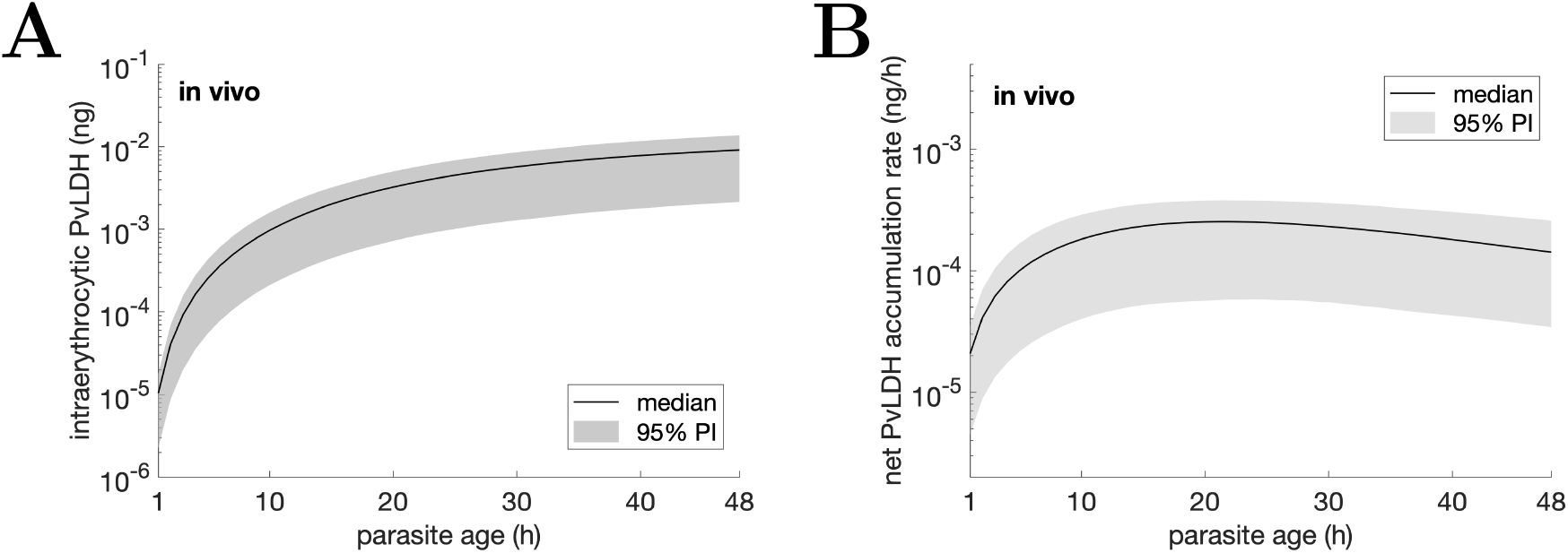
Estimated *in vivo* intraerythrocytic PvLDH level and the net accumulation rate of intraerythrocytic PvLDH over an in vivo asexual life cycle. The net intraerythrocytic PvLDH accumulation rate shown in B is the rate-of-change (the derivative) of the intraerythrocytic PvLDH shown in A. The curve and shaded area represent the median and 95% prediction interval (PI) of model predictions.

**Figure 5:**
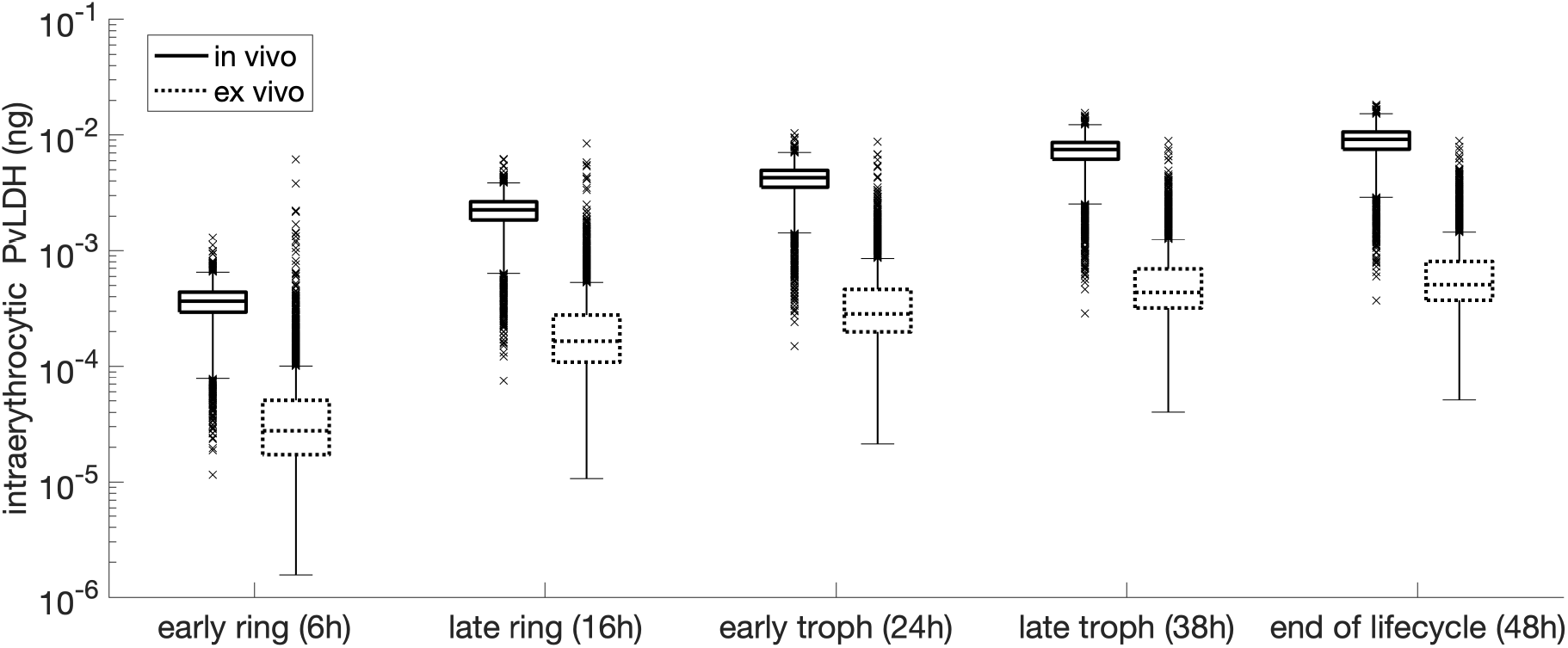
Comparisons between estimated *in vivo* and *ex vivo* intraerythrocytic PvLDH levels at each developmental stage in the asexual life cycle; early ring (6h post-infection), late ring (16h), early trophozoite (24h), late trophozoite (38h) and end of life cycle schizonts (48h). Note that the extended length of *ex vivo* life cycle (72h) was normalised to 48h (i.e., 72h/1.5 = 48h) to facilitate comparisons.

Based on the marginal posterior distribution of the population mean of the *in vivo* decay rate (*λ* in Fig. S3 in the Supplementary Information), we estimate the *in vivo* PvLDH decay rate in the within-host model to have a median of 0.0316 h^-1^ (equivalent to a half-life of 21.9 h) and a 95% CrI of 0.0232—0.0416 h^-1^ (16.7—29.9 h), approximately three times higher than the median estimate of the *ex vivo* PvLDH decay rate (i.e., 0.0106 h^-1^). This suggests that *in vivo* PvLDH clearance from the blood in VIS was three times faster than clearance from the media in *ex vivo* cultures.

## Discussion

In this study, we modelled *ex vivo* and *in vivo* experimental data of parasitemia and PvLDH and estimated key biological parameters that characterise the *ex vivo* and *in vivo* dynamics of PvLDH using Bayesian hierarchical inference. We derived two important parameters, the intraerythrocytic PvLDH level per pRBC (a function of parasite age) and the PvLDH decay rate. We found that *in vivo* and *ex vivo* estimates differed significantly, with intraerythrocytic PvLDH mass estimates *in vivo* at least ten-fold higher across the whole asexual life cycle than those estimated in the *ex vivo* system. At the end of the life cycle, the median intraerythrocytic PvLDH mass estimated by the *in vivo* and *ex vivo* models was 9.4×10^−3^ and 5.1×10^−4^ ng per pRBC, respectively. The rate of intraerythrocytic PvLDH accumulation was fastest during the ring stage in both systems. Similarly, *ex vivo* and *in vivo* estimates of PvLDH decay rates were distinct, with PvLDH clearance from the blood three times faster than the decay rate in culture media. To our knowledge, this is the first study characterising both *ex vivo* and *in vivo* PvLDH dynamics under a single modelling framework and provides an important foundation to advance our understanding of *P. vivax* biology and the development of PvLDH-based diagnostic tools.

The finding that *in vivo* estimates of intraerythrocytic PvLDH mass were at least ten-fold higher than *ex vivo* estimates was unexpected. Growing *P. vivax* in culture remains a major challenge ^17^ as reflected by the presence of morphologically unhealthy parasites and a prolonged *ex vivo P. vivax* lifecycle in our study. We hypothesise that metabolic activity in our cultured parasites may be impaired, including PvLDH production, which would in part explain some of the differences observed between the *ex vivo* and *in vivo* results. Biological conditions or factors that are present/absent *in vivo* versus *ex vivo* may further modulate the production of intraerythrocytic PvLDH. Whether parasite genetic factors may also play a role in varying PvLDH production rates is unclear (i.e., Indian VIS isolate vs. Indonesian clinical isolates). Similarly, the PvLDH decay rate, which measures the rate of decline in PvLDH in either the peripheral blood (*in vivo*) or the culture media (*ex vivo*), was approximately three times faster *in vivo* than *in vitro*. We hypothesise that specific processes/factors that are present *in vivo* but absent in cell culture, such as protein immune-complex formation, metabolic degradation and renal clearance, may lead to faster *in vivo* clearance of PvLDH from peripheral blood. Apart from providing a quantitative understanding of the biological processes, these results highlight the necessity of using *in vivo* estimates of PvLDH dynamics in any future predictive modelling study.

Our estimates of intraerythrocytic PvLDH level as a function of parasite age provide a comprehensive and robust estimate of PvLDH accumulation inside a pRBC across multiple asexual developmental stages. Our *ex vivo* estimates of the intraerythrocytic PvLDH mass and accumulation rate were qualitatively consistent with earlier findings by Barber et al. who identified a rapid increase in PvLDH production during the first 6—12 hours of the life cycle followed by a slow increase or saturation for the remainder of life cycle ^7^. A quantitative comparison of their results with our data was not possible due to the lack of a demonstrably synchronised parasite culture in the earlier study. Our estimates of the pattern of intraerythrocytic PvLDH accumulation over time are also consistent with transcriptomic data whereby levels of *P. vivax* LDH mRNA peak across the 6h to 24h interval, followed by a gradual decline until the end of the lifecycle ^18^.

This model also provides a platform to further evaluate the utility of *P. vivax* biomarkers for non-invasively quantifying the hidden parasite biomass seen with this infection ^7,8,13,14^. This would require the availability of sufficiently sensitive and accurate assays for measuring PvLDH, and other potential *P. vivax* specific biomarkers, in plasma as well as whole blood. Serial measurement of plasma and whole blood PvLDH and parasitemia in patients with symptomatic malaria infection as well as in patients with asymptomatic sub-patent infection would provide additional key data to improve the accuracy of the model.

Our study has several limitations. Our mathematical model does not account for the presence of gametocytes. This is unlikely to have a material impact on the inferred parameters governing the asexual replication cycle as the sexual commitment rate per cycle is less than 1% ^19^. Secondly, we assumed that the intraerythrocytic PvLDH could only be released to plasma upon rupture of viable or dead pRBC. This assumed mechanism of PvLDH release was sufficient to reproduce the *ex vivo* experimental observation, but we cannot exclude gradual release and there have been studies showing that *P. falciparum*-specific LDH can be released from intact pRBC by extracellular vesicles ^20^. This has not been reported for *P. vivax*.

In conclusion, we have established a modelling framework for characterising PvLDH dynamics and provided *ex vivo* and *in vivo* estimates of both intraerythrocytic PvLDH level (including PvLDH levels for each developmental stage in the *P. vivax* asexual life cycle) and PvLDH decay rate. Our work significantly advances our quantitative understanding of PvLDH dynamics and pave the way for the better understanding of *P. vivax* pathobiology and the development of novel methods/tools based on the biomarker to support diagnosis and control of *P. vivax* infections.

## Materials and Methods

### *Plasmodium vivax* culture and microscopy

The *ex vivo* intraerythrocytic PvLDH level was estimated based on experimental data from *ex vivo P. vivax* cultures. Parasite isolates were obtained from malaria patients with *P. vivax* mono-infection as part of a larger surveillance study in Timika, Papua, Indonesia. Written informed consent was obtained for collection of peripheral venous blood. The work was approved by the Human Research Ethics Committees of Gadjah Mada University, Indonesia (KE/FK/0505/EC/2019), and Menzies School of Health Research, Australia (HREC 10-1937). Data from two experiments with optimal cultures, starting parasitemia (>1%) and initial staging (>90% young rings) were selected for the modelling work to satisfy the compatibility between the data and our mathematical model and the inclusion of all developmental stages over the asexual lifecycle.

Fresh *P. vivax* field isolates were cultured at 10% haematocrit in warmed McCoy’s 5A media supplemented with HEPES, gentamycin, D-glucose, L-glutamine and human serum, as previously described ^4,21^. For each experiment, 500 μL of culture suspension was added into each well of a 48-well plate and incubated in a candle jar at 37°C within 2 hours of blood collection. The entire volume of two wells were sampled every 6 hours, resulting in a total of 10 timepoints being collected over a 54-hour culture period. Exact timepoints were recorded to the closest half hour. At each timepoint, suspensions from the two duplicate wells were centrifuged at low speed to gently pellet the red cells. Culture media and red cell pellets were separated and immediately frozen at -80°C. Thick and thin blood smears were prepared for each well at each timepoint and stained with 3% Giemsa solution for 45 minutes at room temperature. Smears were examined by a WHO-certified expert microscopist. Parasite levels were calculated in the thin smear from the number of parasites seen in at least 1000 red cells and converted to total parasite count per well using the average number of red cells per well 6.44×10^8^ estimated based on automated red cell counts and haematocrit data from 19 patients with uncomplicated vivax malaria in a previous cohort ^22^. Healthy parasites were morphologically staged into rings, trophozoites, schizonts and gametocytes, and the fractions of ring, trophozoite and schizont stages are shown in Fig. 1 for the purpose of determining the length of *in vitro* life cycle.

### Inhouse PvLDH ELISA

The concentration of PvLDH was measured in frozen culture media and RBC pellets using an inhouse ELISA as previously described ^7^. RBC pellets were freeze-thawed four times prior to the assay. ELISA plates were read on a GloMax plate reader (Promega, Wisconsin, US). Data were converted from concentration to total mass based on the volume of media/pellet. PvLDH in the media was a measure of PvLDH released by the parasites, while PvLDH in the pellets was a measure of intracellular PvLDH produced by the parasites.

### *In vitro* PvLDH decay

The *in vitro* PvLDH decay rate was estimated by fitting an exponential decay model (Eq. 1 below) to the *in vitro* measurements of human PvLDH decay over time. Human PvLDH from a plasma sample of known PvLDH concentration was introduced to a 10%-hematocrit suspension of healthy uninfected RBCs and cultured in a 48-well plate without parasites under the same conditions described above. Culture media was collected from duplicate wells and frozen at time zero, 30 minutes, then every 6-8 hours over a total of 54 hours. PvLDH concentration in the media at each timepoint was measured by inhouse ELISA as described above and presented in nanograms per mL (Fig. S4 in the Supplementary Information). The exponential decay model is given by

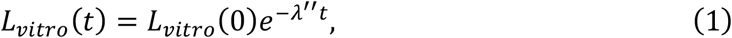

where *L*_*vitro*_*(t)* represents the *in vitro* PvLDH level at time *t* since the first measurement and *λ*^″^ is the *in vitro* PvLDH decay rate (note that we use the double prime superscript to differentiate it from the *ex vivo* and *in vivo* decay rates introduced later).

Model fitting was conducted in Bayesian framework and samples of the target posterior distribution were generated using Hamiltonian Monte Carlo, The prior distributions for *L*_*vitro*_(0) and *λ*^″^ were uniform distributions U(500, 1000) and U(0, 0.1), respectively. Four Markov chains with starting points randomly chosen from the prior distributions were generated and each chain returned (after a burn-in period of 1000 samples) 1000 samples which were used to estimate model parameters and produce marginal posterior distributions. Model fitting was implemented in R (version 4.0.5) and Stan (version 2.21.5). Data and computer code are publicly available at https://doi.org/10.26188/23256413.v1.

Posterior predictive checks of the model fitting and the marginal posterior distributions of *L*_*vitro*_(0) (initial PvLDH) and *λ*^″^ are shown in Figs. S4 and S5, respectively. The posterior samples of *λ*^″^ can be fitted by a normal distribution with a mean of 0.0106 and a standard deviation of 4.1565×10^−4^. This normal distribution is used as the prior distribution for the *ex vivo* decay rate of PvLDH in the *ex vivo* model which is introduced in the next section.

### *Ex vivo* mathematical model of *P. vivax* and PvLDH dynamics

We developed a mathematical model of *P. vivax* and PvLDH dynamics in the *ex vivo* culture system (referred to as the *ex vivo* model) which was fitted to the *ex vivo* experimental data. The parasite maturation dynamics in the *ex vivo* model is structurally similar to the within-host model (described below) except that there is no parasite replication due to the lack of reticulocytes in the culture:

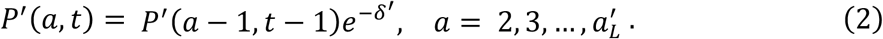

We use *P*′ (*a, t*) (with a prime) to represent the parasite number in the culture and *P*′ (1, *t*) = 0 for all *t* > 0. Since we will introduce an *in vivo* model (called the within-host model) later and the two models share several compartments and parameters that represent the same biological quantities or processes but differ in environment (i.e., *ex vivo* vs. *in vivo*), for those shared compartments and parameters, we use letters with a prime to indicate *ex vivo* quantities. For example, 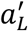 is the length of *ex vivo* asexual life cycle and may differ from its *in vivo* counterpart *a*_*L*_ due to a potential environment-induced variation, which was evidenced in *P. falciparum* ^23^. Since in the *ex vivo* experiment the parasites taken from clinical patients may contain a small amount of schizonts that are not fully mature or ruptured, we chose the initial age distribution of the parasites in the culture *P*′ (*a*, 0) to be

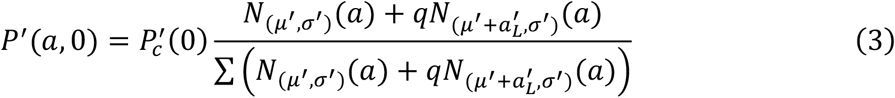

where *N*_(_*μ*′, *σ*′ _)_*(a)* and 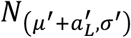 are the probability density functions of the normal distribution *N*(*μ*′, *σ*′) and 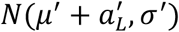 truncated between 0 and 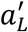 and binned every one hour, representing the age distributions of parasites in the current and the preceding replication cycles, respectively. Since the number of parasites in the current cycle is larger than that in the preceding cycle because of parasite replication, we introduce a parameter *q* ∈ [0,1] to capture this difference (note that *q* can be considered as the reciprocal of the parasite multiplication factor). The probability density is normalised to the sum of the probability density (denominator of Eq. 3) such that the sum of *P*′ (*a*, 0) over all ages is equal to the initial parasite number 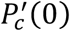 (which will be explained later by Eq. 7). The dead cells that have not ruptured in the culture follow:

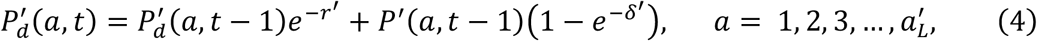

where *r*′ is the *in vitro* rate of dead cell rupture.

The *ex vivo* experiments measured PvLDH in two compartments, PvLDH in the pellet (intracellular PvLDH) and in the media (extracellular PvLDH). The dynamics of PvLDH in the media (denoted by *L*_m_) are modelled by the following difference equations:

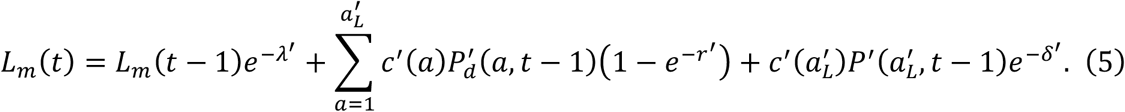

The first term on the righthand side of Eq. 5 represents a reduced amount of PvLDH compared to the amount at the preceding time *L*_m_(*t* − 1) due to natural degradation at rate *λ*′. The second term represents PvLDH released from the ruptured dead cells. The third term represents the amount of PvLDH released into plasma from the rupturing pRBCs. *c*′*(a)* is the cumulative mass of PvLDH inside a single pRBC (note that the cumulative mass is a net result of PvLDH production and degradation inside the pRBC). Since it was found that high LDH activity in the regulation of the formation of pyruvate was associated with mature parasites (i.e., trophozoites and schizonts) ^4^, *c*′ *(a)* is heuristically modelled by an increasing sigmoidal function of parasite age *a*:

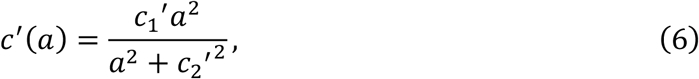

where *c*_1_′ and *c*_2_′ are the tuning parameters modulating the shape of the sigmoidal curve. The initial PvLDH level in culture media *L*_*m*0_ = *L*_*m*_(0) is also a model parameter. The net accumulation rate of intraerythrocytic PvLDH is the derivative of *c*′ *(a)* with respect to *a* and is given by 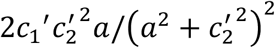. Model parameters and associated information including prior distribution for the Bayesian inference (see below) are provided in Table 1. Note that the length of *ex vivo* life cycle is chosen to be 72h based on that the *ex vivo* life cycle was estimated to be approximately 1.5 times of the *in vivo* life cycle which is 48h. In detail, it was observed in the lower panels of Fig. 1 that the *ex vivo* duration of trophozoite stage, which is approximately the time between 5h culture time (which is followed by a significant drop in the ring fraction) and 41h culture time (which is followed by a significant drop in the trophozoite fraction), is 36h. Since the duration of *in vivo* trophozoite stage is approximately 24h (19h to 42h post-invasion in a 48h life cycle), the ratio of *ex vivo* trophozoite duration to *in vivo* trophozoite duration is 1.5, which, if applied to all other stages (i.e., ring and schizont), results in an estimated *ex vivo* lifecycle of 72h.

Table 1: Parameters of the *ex vivo* model. The prior distribution for the population mean of each model parameter is a uniform distribution with biologically plausible boundaries specified in the table. Note that the *ex vivo* length of asexual life cycle 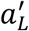 was fixed to be 72h, as explained in the main text. The prior for *λ*^+^ was determined based on a modelling study of an independent *in vitro* experiment where decay data of a human PvLDH were collected and fitted by an exponential decay model (Eq. 1; see the first section of Materials and Methods for details). We assigned an upper limit of *L*_m0=_ of 1.755ng because the initial PvLDH level in culture media was below the limit of quantification of 3.9ng/mL, which is equivalent to 1.755ng based on the culture media volume of 0.45mL. Priors for other parameters were set to be uniform distributions with bounds selected either based on biological plausibility (e.g., 0 as a lower bound) or to be some appropriate values to avoid any resolvable truncation of the posterior distribution (e.g., those upper bounds shown in the table).

Table 1: Parameters of the *ex vivo* model.
| Parameters of the *ex vivo* model |  |  |  |
| --- | --- | --- | --- |
| Parameter | Unit | Description | Prior distribution |
| $\mu'$ | h | mean of the normal distribution defining the initial parasite age distribution | U(0, 24) |
| $\sigma'$ | h | SD of the normal distribution defining the initial parasite age distribution | U(0, 20) |
| $q$ | (unitless) | a fraction parameter | U(0, 1) |
| $c_1'$ | ng | maximum level of intraerythrocytic PvLDH | U(0, 0.01) |
| $c_2'$ | h | parasite age where the intraerythrocytic PvLDH level reaches $c_1/2$ | U(0, 72) |
| $\delta'$ | $h^{-1}$ | *ex vivo* death rate of parasites | U(0, 0.5) |
| $r'$ | $h^{-1}$ | *ex vivo* rupture rate of dead parasite | U(0, 1) |
| $\lambda'$ | $h^{-1}$ | *ex vivo* decay rate of PvLDH | N(0.0106, $4.1565 \times 10^{-4}$)<br>within (0, 0.1) |
| $L_{m0}$ | ng | initial level of PvLDH in culture media | U(0, 1.755) |
| $a_L'$ | h | *ex vivo* length of life cycle | Fixed to be 72 |

The *ex vivo* model can generate the following experimentally measured quantities that are fitted to the *ex vivo* data:

- The number of parasites in the culture (denoted by 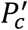) is given by the sum of *P*′ (*a, t*) over all ages:

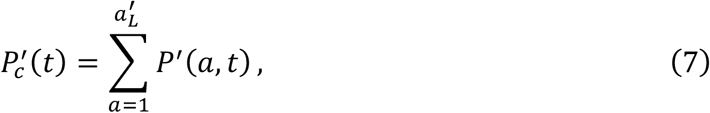
- The PvLDH in the pellet (denoted by *L*_*pe*_) is given by

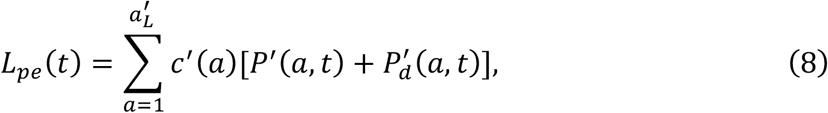

- Total PvLDH in the media is given by *L*_m_(*t*), which is obtained by solving Eq. 5.

To predict the fractions of parasites in different stages (i.e., the predicted fractions of rings, trophozoites and schizonts in Fig. 1), we derive the mathematical expressions of those fractions using the *ex vivo* model:

- The fraction of rings (defined to be fraction of the parasites in the age range of 1—27h based on the prolonged *ex vivo* life cycle of estimated 72h):

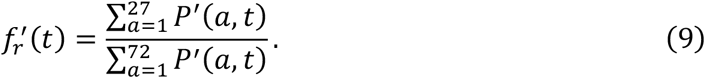

- The fraction of trophozoites (defined to be fraction of the parasites in the age range of 28—63h based on the prolonged *ex vivo* life cycle of estimated 72h):

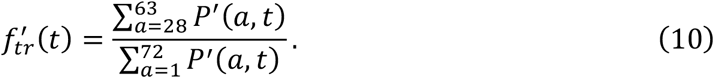

- The fraction of schizonts (defined to be fraction of the parasites in the age range of 64— 72h based on the prolonged *ex vivo* life cycle of estimated 72h):

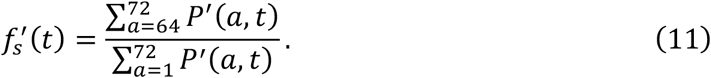

### Volunteer infection study (VIS)

The characterisation of *in vivo* PvLDH dynamics was based on longitudinal parasitemia and PvLDH concentrations in eight malaria-naïve healthy male and female (non-pregnant, non-lactating) subjects who participated in a Phase 1b *P. vivax* induced blood-stage malaria clinical trial ^24^ conducted by the clinical unit Q-Pharm Pty Ltd at QIMR Berghofer Medical Research Institute between 2016 and 2017. The study was approved by the QIMR Berghofer Medical Research Institute Human Research Ethics Committee and the Australian Defense Human Research Ethics Committee. Study design, procedures and main results have been published elsewhere ^24^. Briefly, subjects were inoculated intravenously on day 0 with approximately 564 viable *P. vivax-*infected RBCs originally isolated from a patient with *P. vivax* malaria acquired in India. Parasitemia was measured by 18S qPCR ^25^ and monitored with twice daily blood sampling from day 4 until treatment on days 8-10, and then more frequently until 20 days after treatment. Subjects were treated with a standard course of oral chloroquine (CQ) over 3 days on day 9 (n=1) or day 10 (n=7), as described in ^24^.

### Quansys 5-plex ELISA

PvLDH levels in VIS were measured in whole blood samples collected between day 4 and day of treatment, and every 12 hours after treatment until 120 hours. PvLDH levels in the samples above were measured using the Q-Plex™ Human Malaria assay (Quansys Biosciences, Logan, UT, USA) as previously described ^26^. The Q-View™ Imager Pro was used to capture chemiluminescent images of each plate which was then quantitatively analysed using Q-View™ software (Quansys Biosciences).

### Within-host model of *P. vivax* and PvLDH dynamics for VIS data

We developed a within-host model to capture *in vivo P. vivax* and PvLDH dynamics for the VIS. We adopted the difference equation model structure previously developed ^19,27–29^ to model parasite replication dynamics. In detail, let *P*(*a, t*) represent the number of pRBC in the peripheral blood and depends on both parasite age *a* and time *t* (both in unit of hours), we have:

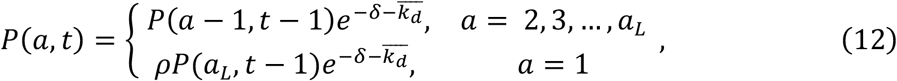

where *a*_*L*_ is the *in vivo* length of *P. vivax* asexual replication cycle and *t* takes integer hours (i.e., *t* = 0, 1, 2, 3, …). *δ* is the *in vivo* death rate of pRBCs. *ρ* is the schizont-to-ring expansion factor indicating the average number of new rings formed due to the rupture of a single schizont from the previous replication cycle. Note that *ρ* is different from the parasite multiplication factor (PMF) or parasite multiplication rate (PMR), which indicates the average amplification rate of parasite number over one asexual life cycle ^30^, and the relationship between *ρ* and PMF follows 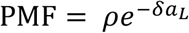. Note that the PMF will be less than the schizont-to-ring expansion factor unless the parasite death rate *δ* is zero. All survival parasites with age *a*_*L*_ will rupture at the next time step. 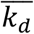 is the average rate of drug-induced parasite killing during each time step (i.e., one hour) and is given by the average of instantaneous killing rate *k*_8_(*t*) at the boundaries of the time step from *t* − 1 to *t*, i.e., 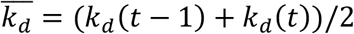 where

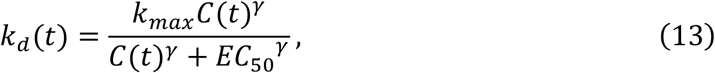

*k*_max_ is the maximum rate of parasite killing by CQ and *EC*_50_ is the half-maximal effective concentration at which the killing rate reaches 50% of the maximum killing rate. *γ* is the Hill coefficient determining the curvature of the dose-response curve. *C*(*t*) represents the effective CQ concentration in the central compartment at time *t* and a CQ concentration versus time profile for each volunteer was simulated using the pharmacokinetic model of CQ developed by Abd Rahman et al. ^31^. Parameter values used in the simulation are provided in Table 2 (the plasma samples column) of Abd Rahman et al.’s article.

The dynamics of dead pRBCs at age *a* and time *t, P*_8_(*a, t*), are modelled by:

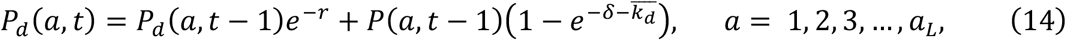

where *r* is the rate of dead pRBC clearance from the peripheral blood.

We assume the age distribution of the 564 pRBCs at *t* = 0 follows a normal distribution *N*(*μ, σ*) truncated between 0 and *a*_-_ and binned by one hour, and parasites with age *a* are those in the bin (*a* − 1, *a*). *μ* and *σ* are the mean and standard deviation of the normal distribution respectively.

For the PvLDH turnover dynamics, the mass of PvLDH (in ng) in the plasma, denoted by *L*_p_(*t*), increases due to the rupture of fully mature pRBCs (schizonts at the end of asexual replication cycle) to the peripheral blood and decreases due to natural degradation. The discrete model for *L*_p_(*t*) is given by:

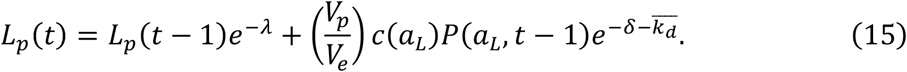

The PvLDH concentration in the plasma at *t* = 0, *L*(0), was set to be zero (i.e., no plasma PvLDH before the inoculation with parasites). The first term on the righthand side of Eq. 15 represents a reduced amount of PvLDH compared to the amount at the preceding time *L*(*t* − 1) due to natural degradation at rate *λ*. The second term represents the amount of PvLDH released from the ruptured pRBCs to the plasma. *V*_p_ and *V*_e_ represent the volume of plasma and the volume of extracellular fluid (both in mL) and are given by:

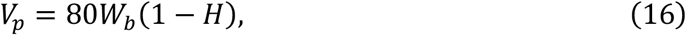

and

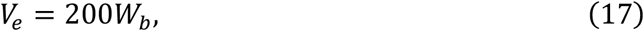

where *W*_*b*_ is body weight and *H* is haematocrit ^10,12^. Note that we assume dead cells do not contribute to the plasma in the peripheral blood because they are quickly phagocytosed by macrophages and the enclosed PvLDH is assumed to remain inside the macrophages. Similar to Eq. 6, the cumulative intraerythrocytic PvLDH level *c(a)* for *in vivo* parasites is given by:

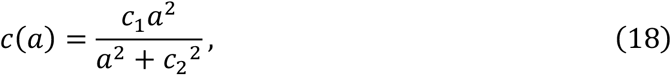

where *c*_1_ and *c*_2_ are the tuning parameters modulating the shape of the sigmoidal curve. The net accumulation rate of intraerythrocytic PvLDH is the derivative of *c(a)* with respect to *a* and is given by 2*c*_1_*c*_2_^2^*a*/(*a*^2^ + *c*_2_^2^)^2^.

The mass of whole blood PvLDH (denoted by *L*_*h*_(*t*)) is given by the sum of plasma PvLDH *L*_*p*_(*t*) and the mass of PvLDH inside either viable parasites or dead pRBCs:

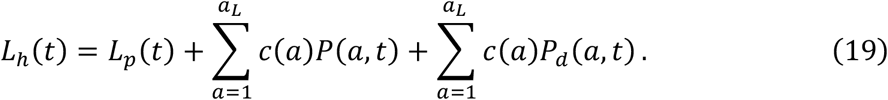

Model parameters and associated information including prior distributions for Bayesian inference are provided in Table 2.

Table 2: Parameters of the within-host model. The prior distributions are chosen for the population means of the model parameters. The lower bounds of the model parameters for those with a prior uniform distribution were selected based on biological plausibility (e.g., 0 as a lower bound). The priors for *c*_1_ and *c*_2_ are chosen to be the posterior distributions obtained from the *ex vivo* model fitting (see the green curves in Fig. S1 in the Supplementary Information). The priors for *k*_*max*_ and *EC*_50_ were chosen based on the estimates from Abd Rahman et al. ^31^.

| Parameters of the within-host model |  |  |  |
| --- | --- | --- | --- |
| Parameter | Unit | Description | Prior distribution |
| $\mu$ | h | mean of the normal distribution defining the initial parasite age distribution | U(0, 48) |
| $\sigma$ | h | SD of the normal distribution defining the initial parasite age distribution | U(0, 80) |
| $\rho$ | (unitless) | schizont-to-ring expansion factor | U(0, 100) |
| $c_1$ | ng | maximum level of intraerythrocytic PvLDH | logN(-7.15, 0.65)<br>within (0, 0.1) |
| $c_2$ | h | parasite age where the intraerythrocytic PvLDH level reaches $c_1/2$ | N(29.17, 8.40)<br>within (0, 72) |
| $\delta$ | h<sup>-1</sup> | *in vivo* death rate of parasites | U(0, 1) |
| $r$ | h<sup>-1</sup> | clearance rate of dead pRBCs from circulation | U(0, 2) |
| $\lambda$ | h<sup>-1</sup> | *in vivo* decay rate of PvLDH | U(0, 0.1) |
| $k_{max}$ | h<sup>-1</sup> | maximum parasite killing rate by CQ | N(0.213, $8.7 \times 10^{-3}$)<br>within (0, 1) |
| $EC_{50}$ | ng/mL | CQ's half-maximal effective concentration | N(15, 2)<br>within (0, 50) |
| $\gamma$ | (unitless) | Hill coefficient of the dose-response curve | Fixed to be 2.5 |
| $a_L$ | H | *in vivo* length of asexual life cycle | Fixed to be 48 |

The within-host model can generate the following experimentally measured quantities that will be fitted to the VIS data:

- The number of circulating pRBCs per mL of peripheral blood (i.e., the peripheral blood parasitemia, denoted by *P*_c_(*t*)) is given by: 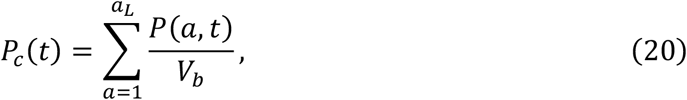

where *V*_*b*_ is the whole blood volume (mL) and is estimated based on 80 mL per kg of the volunteer’s body weight, i.e., *V*_*b*_ = 80*W*_*b*_ ^10^. Note that the initial parasitemia *P*_c0_ = *P*_c_ (0) is fixed to be 564 based on the experimental inoculation size.
- *L*_h_(*t*) was converted from mass to concentration by dividing it by the total blood volume *V*_*b*_, in order to fit to the whole blood PvLDH measurements which are given by concentration.

To predict the fractions of parasites in different stages (i.e., the predicted fractions of rings, trophozoites and schizonts in Fig. 3), similar to what we did with the *ex vivo* model, we derive the mathematical expressions of those fractions using the within-host model:

- The fraction of rings (defined to be fraction of the parasites in the age range of 1—18h based on the *in vivo* life cycle of 48h):

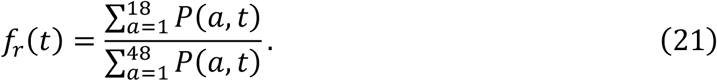

- The fraction of trophozoites (defined to be fraction of the parasites in the age range of 19—42h based on the *in vivo* life cycle of 48h)

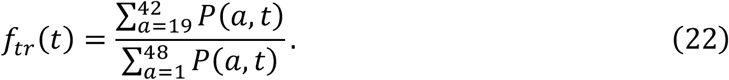

- The fraction of schizonts (defined to be fraction of the parasites in the age range of 43— 48h based on the *in vivo* life cycle of 48h)

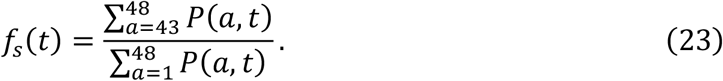

### Relationship between inhouse ELISA PvLDH data and Quansys PvLDH data

The posterior distributions of *c*_1_′ and *c*_2_′ obtained from the *ex vivo* model fitting was used as the prior distributions of *c*_1_ and *c*_2_ in the within-host model. To perform this, we established the relationship between PvLDH concentrations generated using inhouse ELISA and Quansys methods, given that PvLDH concentrations were measured by inhouse ELISA in the *ex vivo* system while concentrations in the VIS were measured by Quansys. Using this relationship, we transformed the VIS PvLDH data to calibrate with the *ex vivo* data measured by inhouse ELISA. The relationship between inhouse ELISA and Quansys PvLDH concentrations was established using a retrospective cross-sectional dataset of 58 vivax malaria patients in Sabah, Malaysia, who had whole blood PvLDH concentrations measured by both inhouse ELISA and Quansys methods ^26^. Using a Bayesian approach, we fitted a linear model to the log-transformed ELISA and Quansys data from Malaysian patients (Fig. S6 in the Supplementary Information). The relationship was estimated to be:

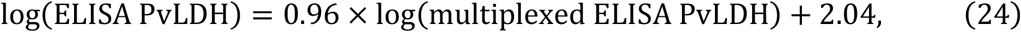

which was used to calibrate the VIS PvLDH concentrations to the *ex vivo* ELISA measurements. The Malaysian patient data and computer code for the model fitting are publicly available at https://doi.org/10.26188/23256413.v1.

### Bayesian hierarchical modelling for the *ex vivo* model and the within-host model

Bayesian hierarchical modelling was performed for model fitting and parameter estimation for the *ex vivo* model and the within-host model, because the experimental data are grouped by patients. The hierarchical model expands the mathematical model (either the *ex vivo* model or the within-host model) by introducing two levels of model parameters:

- *the individual-level parameters* (or simply the individual parameters) are the model parameters applicable to individuals such as patient isolates or volunteers. Each individual owns a set of model parameters that differs from that of other individuals. For example, each of the eight volunteers in VIS has 10 parameters (i.e., those described in Table 2) to determine their own infection kinetics and the 10 parameters for one volunteer differ from the 10 parameters for another volunteer. Therefore, there are in total 80 individual parameters in the hierarchical within-host model.
- *the population-level parameters* (or simply the population parameters) determine how the individual parameters are distributed. We assume individual parameters follow a multivariate normal distribution with a set of means (referred to as the population mean parameters) and standard deviations (referred to as the population SD parameters). Note that since the parameters in our models are positive and bounded due to biological plausibility, we assume that log-transformed individual parameters follow a multivariate normal distribution with log-transformed population means and SDs.

With a hierarchical model (either the *ex vivo* model or the within-host model), fitting the model to experimental data (either the *ex vivo* data or the VIS data) and sampling from the posterior distribution for parameter estimation were implemented in R (version 4.0.5) and Stan (RStan 2.21.5) using the Hamilton Monte Carlo sampling method optimized by the No-U-Turn Sampler ^32,33^. Four chains were randomly initiated, and each chain generated 1,000 (for the ex vivo model fitting) or 2,000 (for the within-host model fitting) samples (excluding burn-in samples), giving in total 4,000 (for the ex vivo model fitting) or 8,000 (for the within-host model fitting) samples drawn from the posterior distribution. Note that the longer chains for within-host model fitting were required to increase effective sample size. The M3 method was used to penalise the likelihood for data below the limit of detection ^34^. Diagnostic outputs, such as 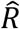 statistic and effective sample size were examined to ensure the convergence of the chains and low sample autocorrelations. The median model prediction and 95% prediction interval (PI) are given by the median and quantiles of 2.5% and 97.5% of the 4,000/8,000 model solutions at each time respectively. For example, to produce the model prediction shown in Fig. 3A, we simulated the within-host model 8,000 times using the 8,000 posterior values of the individual parameters of the within-host model for Volunteer R009 and calculated the median and 95% PI for all time points. The median and 95% credible interval (CrI) of a model parameter (e.g., the PvLDH decay rate) are given by the median and quantiles of 2.5% and 97.5% of the posterior samples of the parameter. To estimate the *in vivo* (or *ex vivo*) intraerythrocytic PvLDH level, posterior samples of the population mean of *c*_1_ and *c*_2_ (or *c*_1_′ and *c*_2_′ for the *ex vivo* model) were put into Eq. 18 (or Eq. 6 for the *ex vivo* model) to obtain the median and 95% PI for all parasite age values (see Fig. 2 for results). Similarly, the *ex vivo*/*in vivo* net PvLDH accumulation rate can be calculated using the same posterior samples of the population mean parameters and the derivatives of Eqs. 6 and 18. All main text and supplementary figures were generated using MATLAB (version 2019b; The MathWorks, Natick, MA). All experimental data, computer code (R and MATLAB) and model fitting results are publicly available at https://doi.org/10.26188/23256413.v1.

## Acknowledgments

We thank Dr Sophie Zaloumis (University of Melbourne) for helpful conversations about the method for model fitting and Dr Louise Marquart (University of Queensland) for her preliminary analysis of the VIS data. We also thank Dr Gonzalo Domingo (PATH) for his contribution to data collection and analysis. The work was supported by the National Health and Medical Research Centre (NHMRC) of Australia: Project Grant (1025319), a Senior Principal Research Fellowship to Nicholas M. Anstey (1135820), Investigator Grants Leadership Level to James McCarthy (GNT2016396) and Julie A Simpson (1196068) and supported in part by the Australian Centre for Research Excellence on Malaria Elimination (1134989). Steven Kho is supported by a Menzies Future Leaders Fellowship.

## Author contributions

NMA, JSM, SB conceived the study; SK, MJG, BEB, KAP, TW, JRP, IKJ conducted the experiments and provided the data; PC, SK, SB analysed the data with contributions from JAS, JMM, NMA, JSM; PC developed the mathematical models and performed model fitting, simulation and analysis; PC and SK wrote the first draft of the manuscript with contributions from all other authors. All authors reviewed and commented on the manuscript.

## Competing interests

We have no competing interests.

## Supplementary information

**Figure S1:**
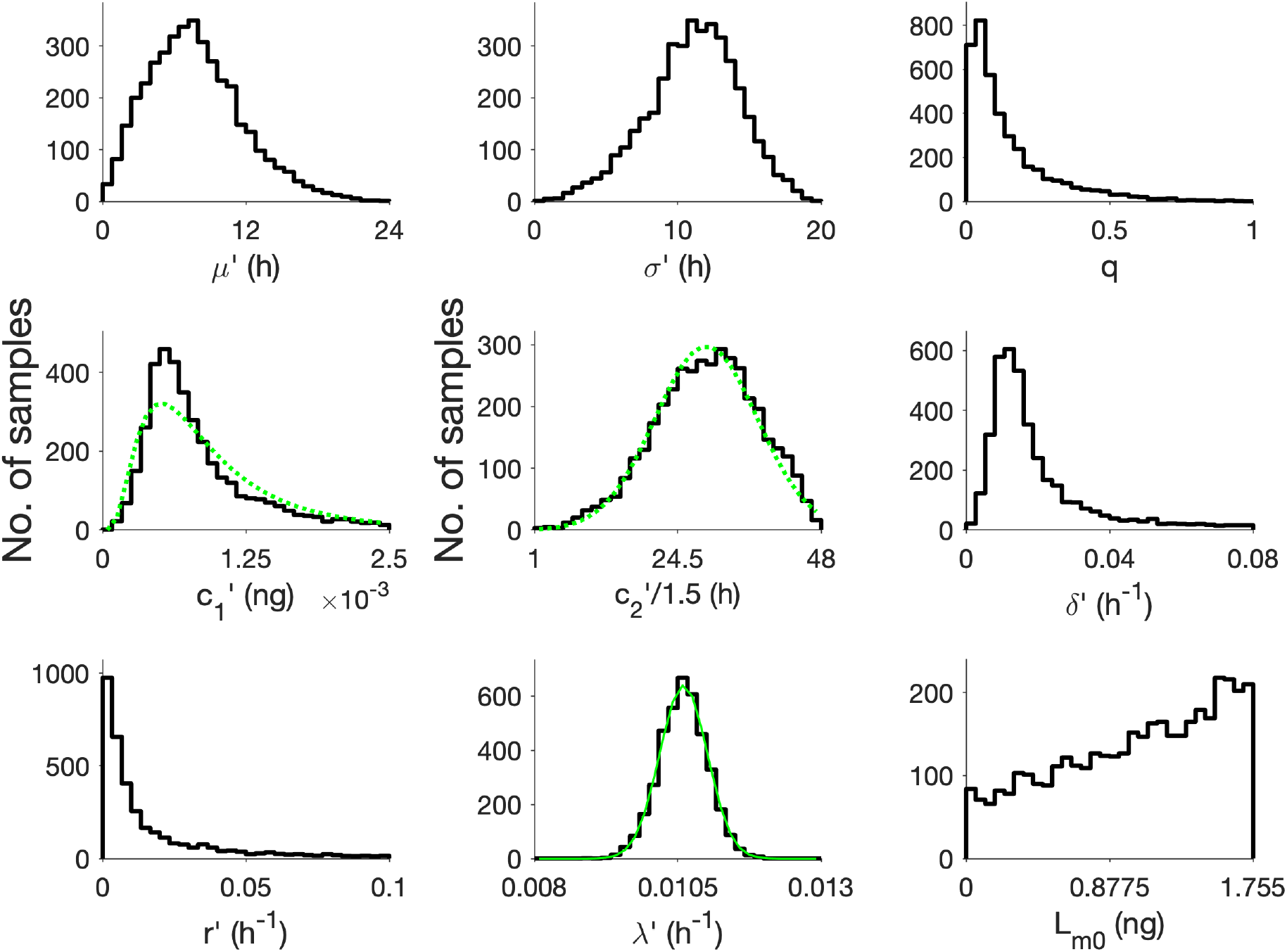
Posterior distributions of the population mean parameters of the ex vivo model. The prior distributions for the model parameters are uniform distributions specified in Table 1 in the main text except that the prior for the *ex vivo* PvLDH decay rate *λ*^+^ is a bounded normal distribution shown by the green curve (and also specified in Table 1). The marginal posterior distributions of *c*_1_′ and *c*_2_′ were empirically fitted by a lognormal distribution logN(−7.15, 0.65) and a normal distribution N(29.17, 8.40), respectively, shown by the green dotted curves. The empirical fits were used as prior distributions for *c*_1_ and *c*_2_ in the within-host model fitting (see Table 2 in the main text). Note that the range of *c*_2_′ was scaled down by 1.5 such that its empirical fit is applicable to the within-host model fitting.

**Figure S2:**
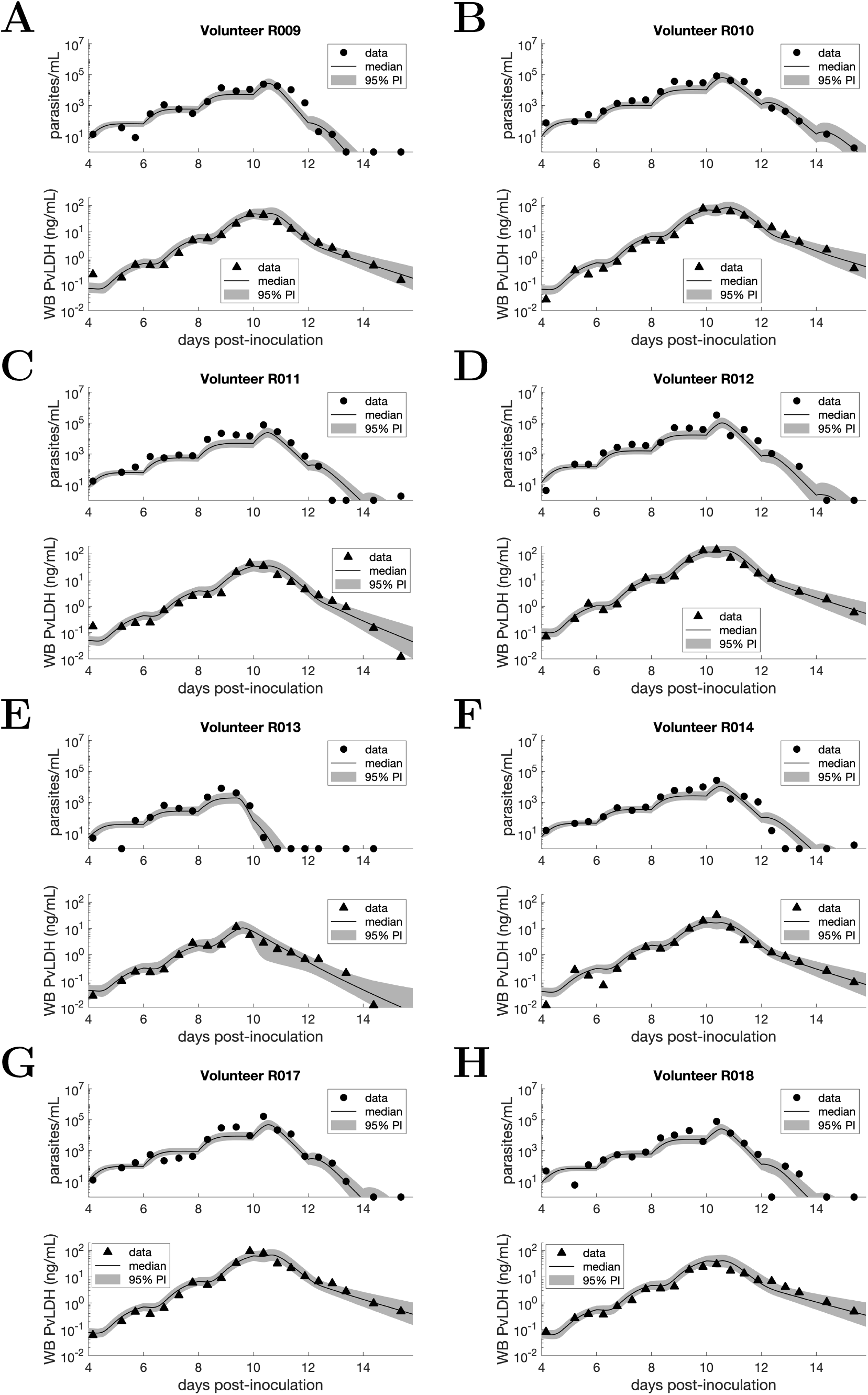
Data and model fits for eight VIS volunteers. Solid dots are clinical data and model fits are shown by the median and 95% PI of the model-predicted distributions of parasitemia and whole blood PvLDH concentration (in ng per mL blood) at different time points.

**Figure S3:**
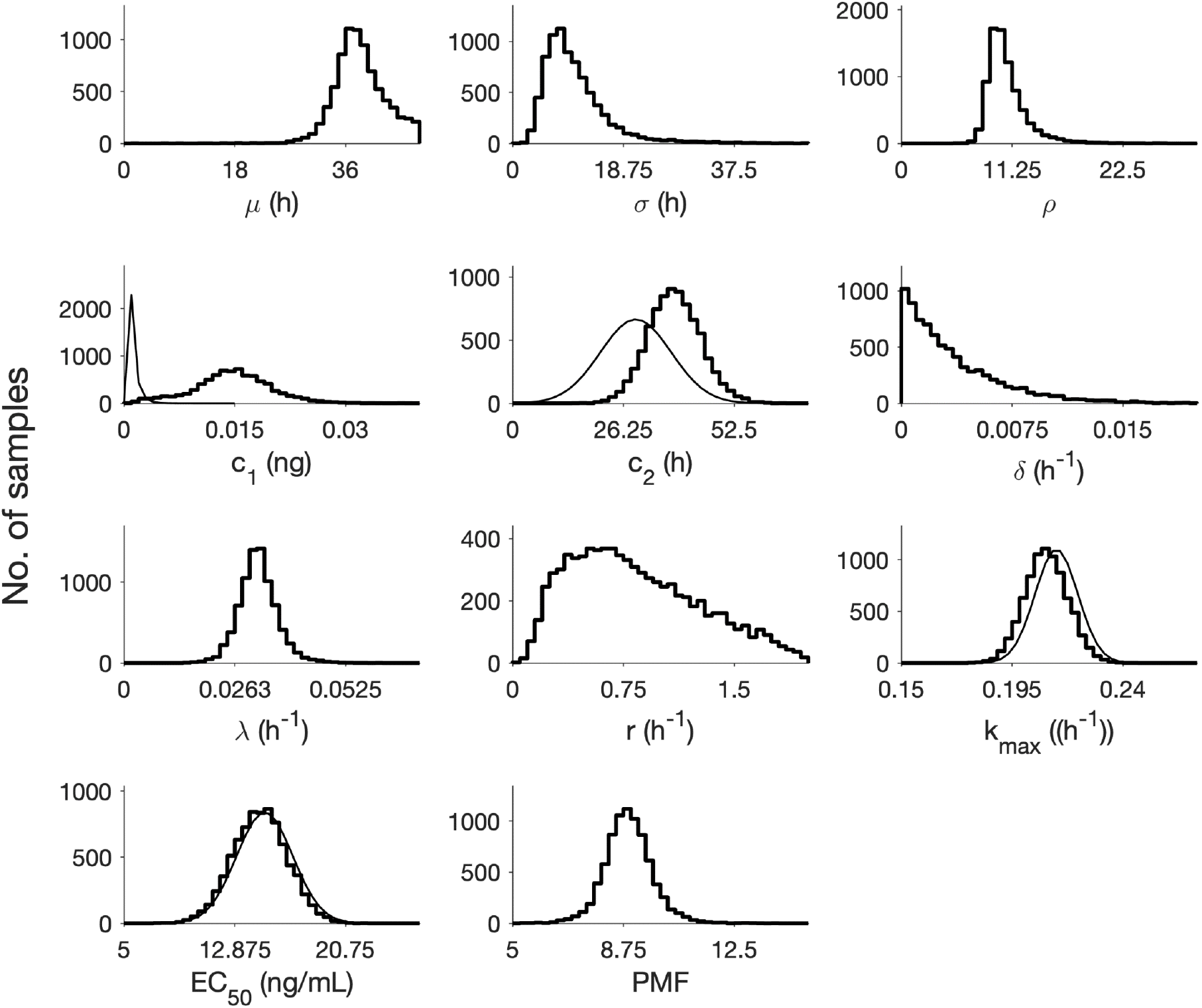
Posterior distributions of the population mean parameters of the within-host model. The prior distributions for the model parameters are either uniform distributions or bounded normal/lognormal distributions, as specified in Table 2. Normal/lognormal priors are shown by the smooth curves. The distribution of the parasite multiplication factor (PMF), which is calculated based on the posterior samples of *ρ* and *δ* and 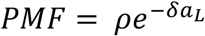, is shown in the last panel.

**Figure S4:**
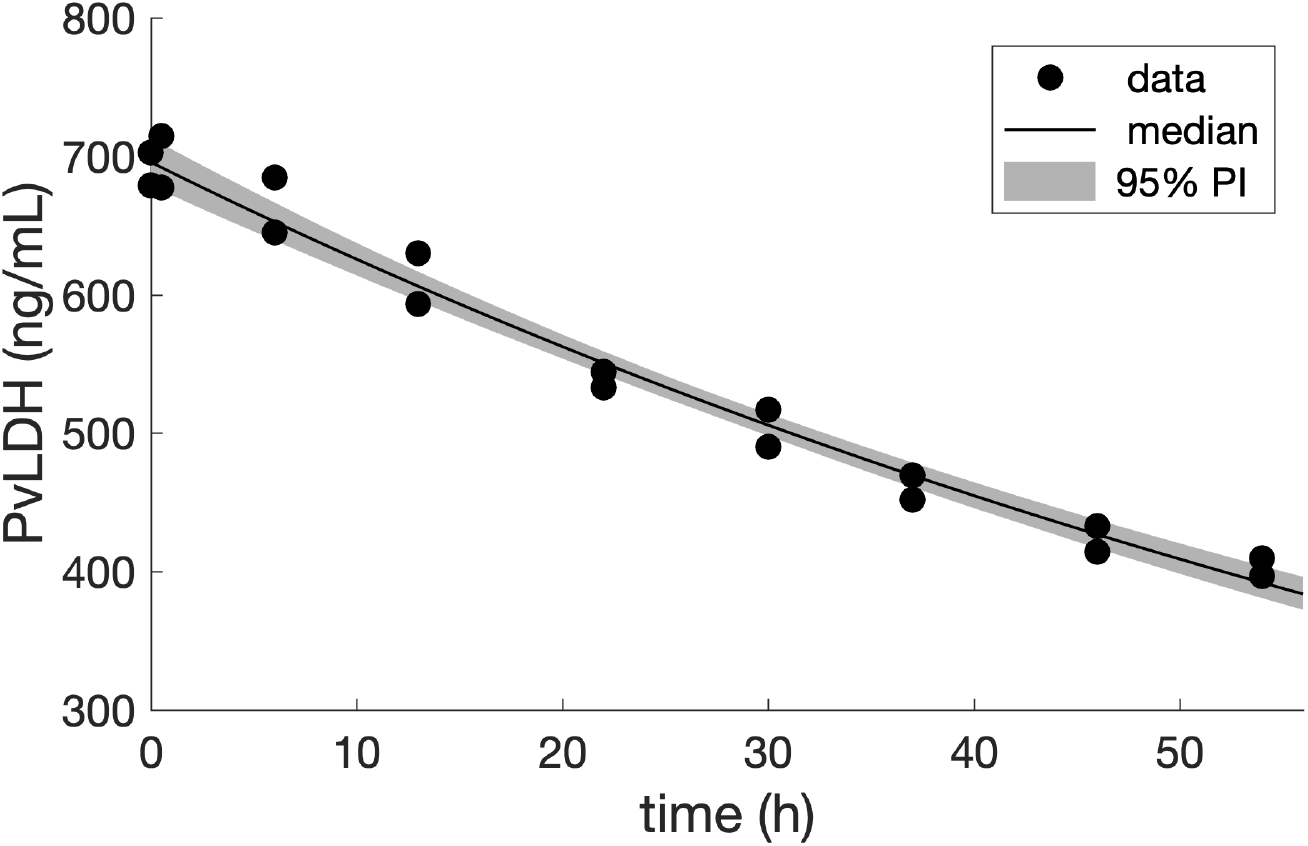
Result of fitting the exponential decay model (Eq. 1 in the main text) to the *in vitro* decay measurements of human PvLDH (see the Materials and Methods for further details). Black dots are experimental data, and experimental measurements were conducted in duplicate at each timepoint. The black curve is the median model prediction based on the 4,000 posterior samples. The shaded region represents the 95% prediction interval whose upper and lower boundaries are the 2.5% and 97.5% quantiles of model predicted PvLDH level at various time points.

**Figure S5:**
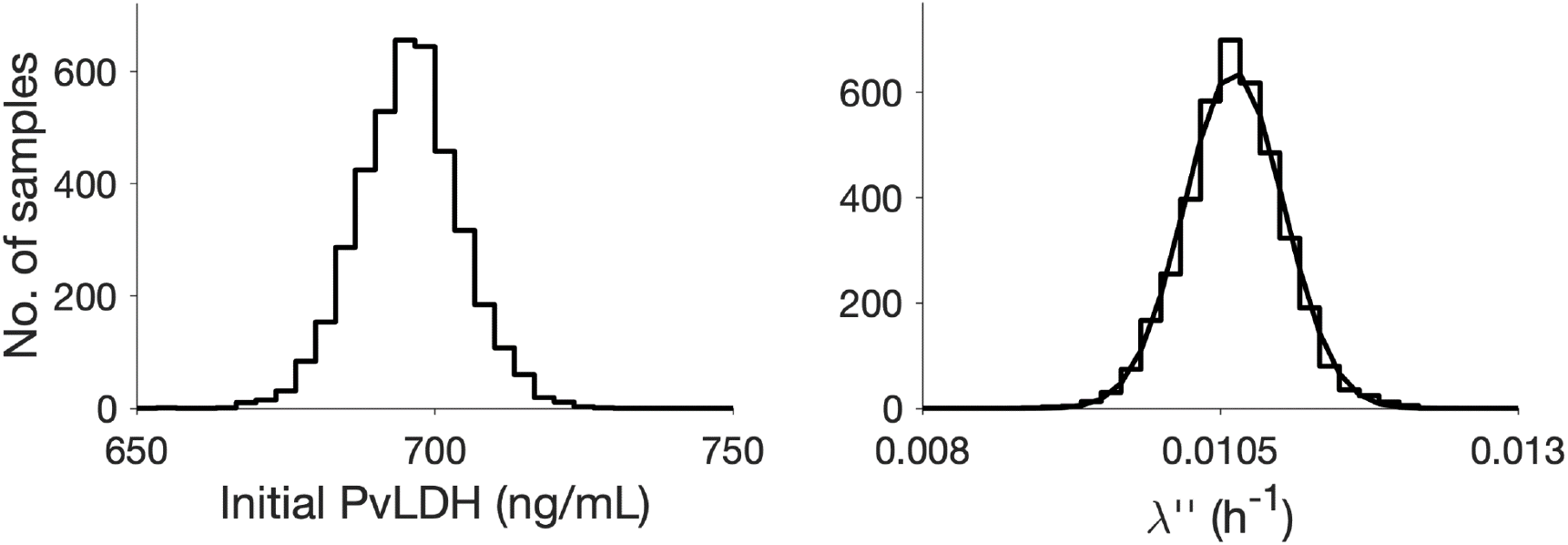
Marginal posterior distributions of *L*_*vitro*_(0) (initial PvLDH) and *λ*″ (which are parameters of the exponential decay model in Eq. 1 in the main text). The prior distributions of *L*_*vitro*_(0) and *λ*″ are uniform distributions U(500, 1000) and U(0, 0.1) respectively. Each of the histograms contains 4000 posterior samples. The histogram of *λ*″ was fitted by a normal distribution N(0.0106, 4.1565×10^−4^) which is shown by the smooth curve. The empirical distribution of *λ*″, N(0.0106, 4.1565×10^−4^), will be used as the prior distribution of the *ex vivo* PvLDH decay rate *λ*″ in the *ex vivo* model fitting.

**Figure S6:**
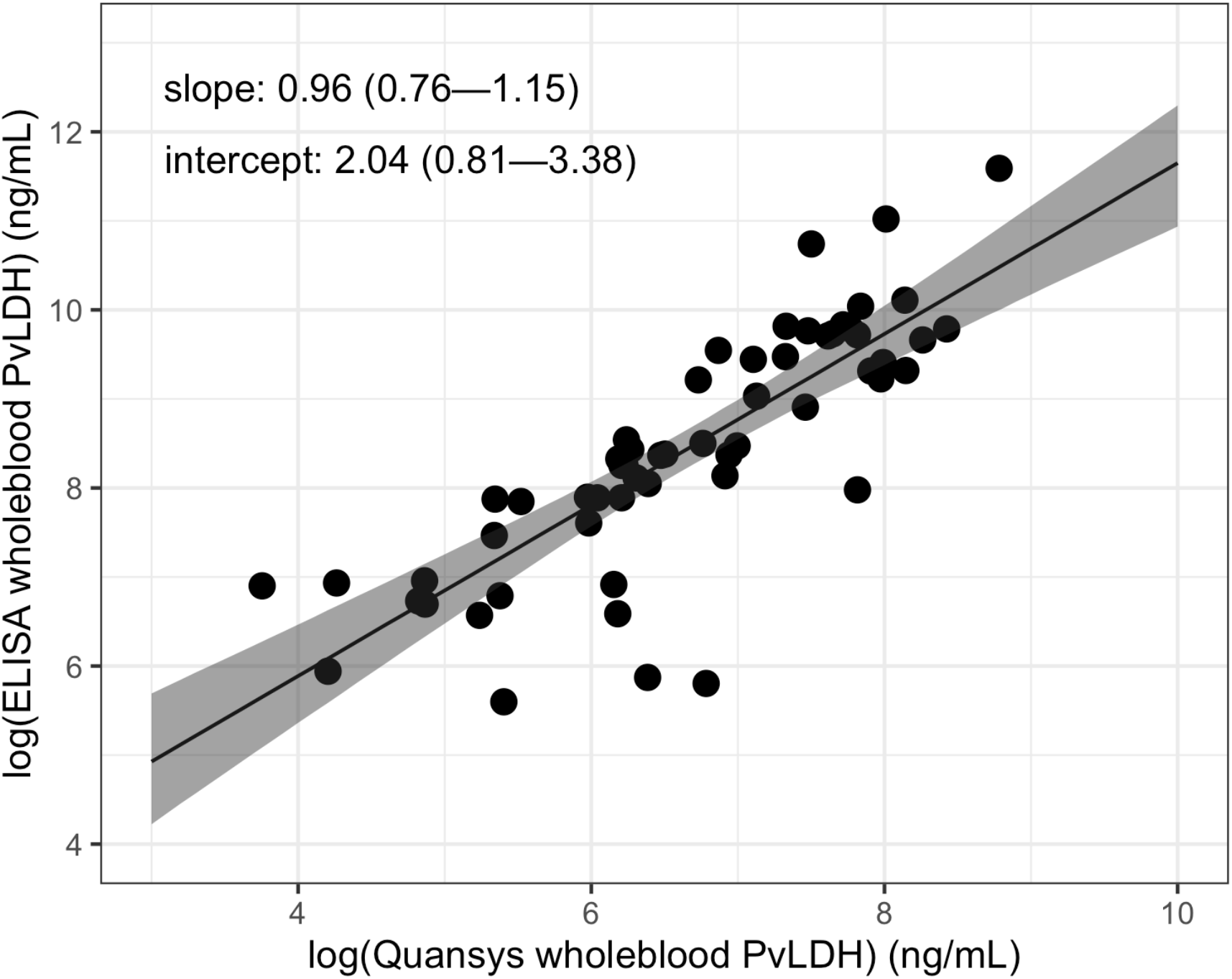
The relationship between Quansys measurement of whole blood PvLDH and inhouse ELISA measurement of whole blood PvLDH. Experimental data (black dots) are obtained from 58 Malaysian patients whose whole blood PvLDH was measured by both methods. The data are log-transformed and fitted by a linear model. The median and 95% prediction interval are shown by the black line and the shaded area, respectively. The median estimates and 95% credible intervals (shown in parentheses) of the slope and intercept parameters of the linear model are also provided in the figure.

## Notes

https://doi.org/10.26188/23256413.v1

### Competing Interest Statement

The authors have declared no competing interest.

